# GABA-glutamate corelease is a mechanism for state-dependent neurotransmission

**DOI:** 10.1101/2025.11.19.689286

**Authors:** Shelley M. Warlow, Dina S. Dowlat, Nick G. Hollon, Lauren Faget, Lucie Oriol, Vivien Zell, Larry S. Zweifel, Thomas S. Hnasko

## Abstract

Ventral tegmental area neurons projecting to lateral habenula (LHb) corelease the main inhibitory and excitatory transmitters, GABA and glutamate (VTA^GG^). Yet the functional role of this synchronous signal remains unclear. We hypothesized that the sign of VTA^GG^ action depends on postsynaptic state in LHb. Ex vivo, activating VTA^GG^ terminals evoked excitatory and inhibitory responses in LHb that varied with postsynaptic membrane potential. In vivo, VTA^GG^ inputs drove net inhibition and supported positive reinforcement that was dependent on GABA, but not glutamate, release. Using chemogenetics to bidirectionally modulate LHb, we found that LHb hyperpolarization shifted VTA^GG^ effects toward excitation, abolishing positive reinforcement, whereas LHb depolarization enhanced net inhibition and positive reinforcement. Thus, the activity state of LHb neurons dictates whether GABA-glutamate corelease is functionally inhibitory or excitatory and can reverse the motivational valence of VTA^GG^ input, supporting a homeostatic role for GABA-glutamate cotransmission with broad implications for disorders of imbalanced motivation.

## Introduction

The ventral tegmental area (VTA) is an important regulator of reward-related motivation. Famous for its dopamine neurons, the VTA is highly heterogenous, containing multiple neuron types including GABA and glutamate neurons, as well as neurons with the capacity to release multiple neurotransmitters^1,2^. For example, a proportion of dopamine neurons express the vesicular glutamate transporter (VGLUT2) enabling glutamate corelease, and a separate population of VGLUT2+ neurons express the vesicular GABA transporter (VGAT) and corelease GABA^3–5^. We and others have shown that optogenetic activation of VTA glutamate neurons and their projections to nucleus accumbens (NAc) promote reinforcement in mice, independent of dopamine co-release^6–9^. A separate population of VTA neurons project to lateral habenula (LHb), and the majority of these express both VGLUT2 and VGAT^4^. The functional role of VTA GABA-glutamate (VTA^GG^) neurons, and specifically why presynaptic neurons would provide synchronized inhibitory and excitatory signals on the same postsynaptic LHb neurons, remains unresolved.

Activity in LHb is positively correlated with aversive events, as well as the expectation of such events occurring in future^10–12^. Direct LHb stimulation drives avoidance behavior, whereas inhibiting LHb promotes place preference and reward-seeking^10,13,14^. These effects are generally attributed to LHb-driven disynaptic inhibition of VTA dopamine neurons^15,16^. Sustained LHb hyperactivity has been linked to depression and drug withdrawal, while LHb hypoactivity may promote manic states or compulsive reward-seeking^17–19^. Accordingly, LHb functions as a hub for stabilizing mood and motivational states.

LHb is also a hotspot for GABA-glutamate cotransmission. In addition to VTA, a majority of LHb afferents from the entopeduncular nucleus and other regions coexpress markers for both GABA and glutamate release^5,20,21^. Across circuits, GABA-glutamate cotransmission has been proposed to support homeostasis, prevent seizures, and facilitate learning^22,23^. Previous works suggest that activation of VGLUT2+ VTA projections to LHb are net inhibitory and sufficient to support self-stimulation, but other assays report aversive effects^4,9,24^, suggesting this input can drive bivalent motivational outcomes. We hypothesized that GABA-glutamate cotransmission homeostatically normalizes postsynaptic activity in LHb: inhibiting when LHb neurons are more active, exciting when they are less active, and thereby producing state-dependent effects on motivated behavior.

## Results

### GABA-glutamate cotransmission in lateral habenula is state-dependent

To selectively target VTA^GG^ neurons we used an intersectional approach to express Channelrhodopsin (ChR2) in dual transgenic VGLUT2-Cre, VGAT-Flp mice (**Fig. 1a-c**). We recorded from LHb neurons while optogenetically stimulating VTA^GG^ terminals. We observed optogenetically evoked excitatory postsynaptic currents (oEPSCs) that were dependent on AMPA-type glutamate receptors, as well as inhibitory postsynaptic currents (oIPSCs) dependent on GABA_A_ receptors (**Fig. 1d-e**). Switching to current clamp revealed that VTA^GG^ stimulation produced either depolarizing or hyperpolarizing optogenetically-evoked postsynaptic potentials (oPSP) across cells (**Fig. 1f**). Crucially, the sign and magnitude of the oPSP varied with resting membrane potential (Vm): more hyperpolarized cells tended to display oEPSPs, whereas more depolarized cells displayed oIPSPs (**Fig 1g-h**). Imposing Vm shifts with current injections confirmed causality: hyperpolarization biased LHb responses to VTA^GG^ stimulation toward excitation, depolarization toward inhibition (**Fig. 1i-j**), with an x-intercept (no net effect) equal to-60 ± 7 mV (**Fig 1k**). This suggests that the net effect of VTA GABA-glutamate cotransmission can fluctuate between excitatory and inhibitory as a function of the activity state of the postsynaptic cell, consistent with a model in which changes in driving force across GABA and glutamate receptors determine the instantaneous sign of VTA^GG^ input, and homeostatically drive Vm toward a midpoint.

**Figure 1.**
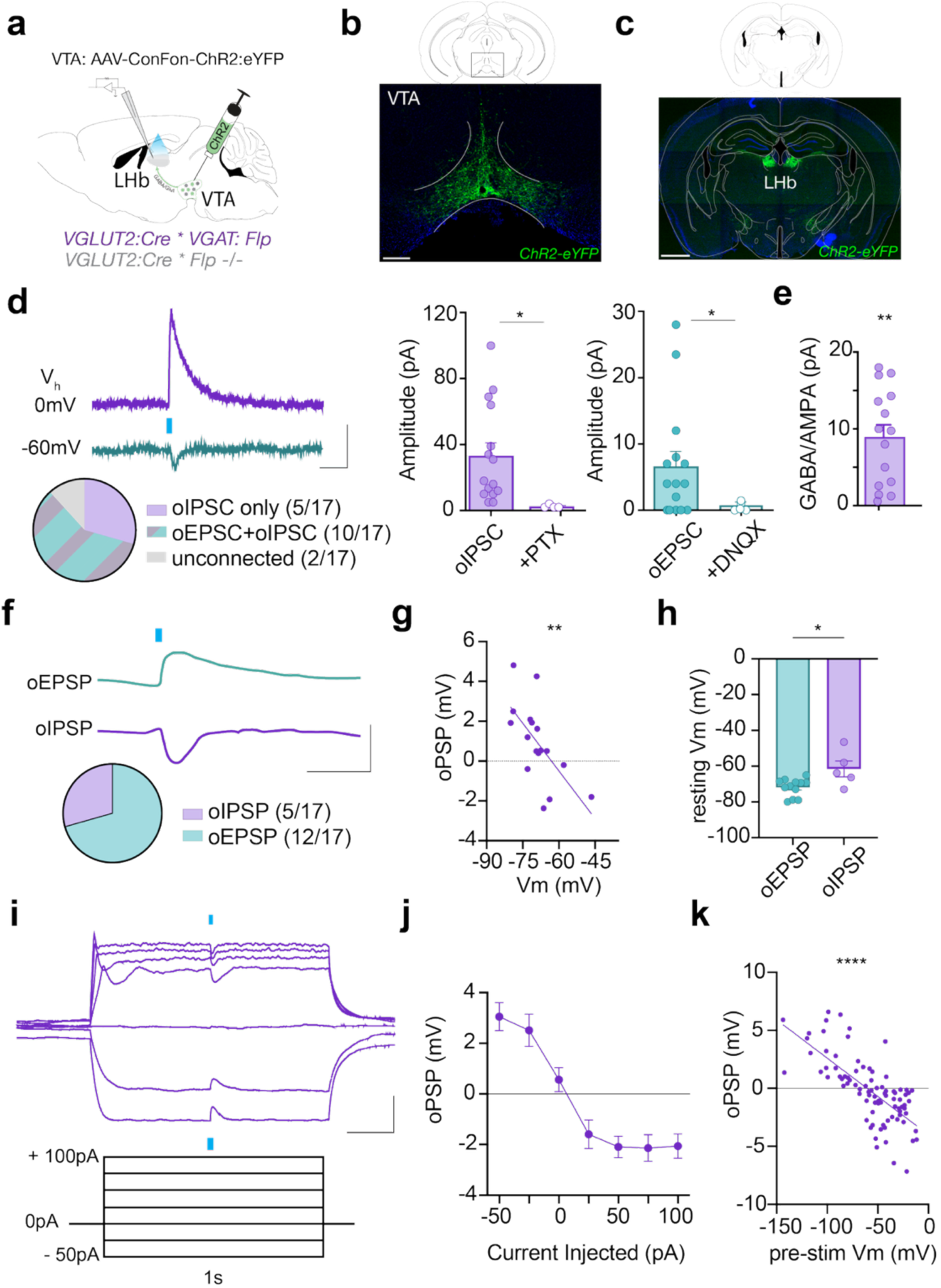
The excitatory versus inhibitory effects of VTA^GG^ input depends on postsynaptic membrane potential in LHb. (A) Strategy to express ChR2:eYFP in VTA neurons positive for both VGLUT2-Cre and VGAT-Flp (VTA^GG^). (B) ChR2 expression in VTA, scale bar = 250 um. (C) Terminal expression of ChR2 in LHb, scale bar = 1 mm. (D) Optogenetically-evoked inhibitory postsynaptic currents (oIPSCs; purple) were observed in LHb neurons in response to 50ms blue light pulses at holding voltage (V_h_) = 0 mV and blocked by picrotoxin (PTX; paired t-test: t_3_=3.4, p=0.04), whereas optogenetically-evoked excitatory postsynaptic currents (oEPSCs; green) were observed at V_h_=-60 mV and sensitive to 6,7-dinitroquinoxaline-2,3-dione (DNQX; paired t-test: t_4_=2.8, p=0.04); scale: x= 50 ms, y=20 pA. (E) Most recorded neurons displayed a GABA/AMPA ratio greater than 1 (one-sample t-test vs. ratio of 1: t_14_=4.9, p=0.0002). (F) In current-clamp mode (I=0), optogenetic-evoked inhibitory postsynaptic potentials (oIPSPs, purple) and-excitatory postsynaptic potentials (oEPSPs, green) were observed in different LHb cells; scale: x=50 ms, y=5 mV. (G) The sign and scale of oPSP displayed as a function of resting membrane potential of postsynaptic LHb cell (Vm; Pearson r =-0.65, p=0.005), and (H) neurons in which terminal stimulation of VTA^GG^ inputs caused an oEPSP were more hyperpolarized compared to cells that responded with an oIPSP (Unpaired t-test: t_15_=2.9, p=0.01). (I) Example trace of a LHb neuron given hyperpolarizing and depolarizing current injections, with blue light pulse delivered halfway through 1s current step; scale: x=100 ms, y=20 mV. (J) Mean oPSPs were positive during negative current steps, and negative during positive current steps (One-way ANOVA: F_6,82_=18.8, p<0.0001). (K) oPSPs plotted by membrane voltage at each corresponding current step (Pearson R correlation: r=-0.71, p<0.0001). Data are shown as mean +/-SEM and/or individual points; *p<0.05, **p<0.01, ****p<0.0001.

### VTA GABA-glutamate cotransmission promotes reinforcement through GABA release

To test the effects of VTA^GG^ projections to LHb on behavior, we intersectionally targeted ChR2 expression to VTA^GG^ neurons and placed fibers bilaterally above LHb to optogenetically stimulate VTA^GG^ terminals (**Fig. 2a-c**). In a two-hole nose-poke task, mice were given the opportunity to poke for optogenetic stimulation (**Fig. 2d**). Mice expressing ChR2 consistently nose-poked to earn optogenetic stimulation, compared to poking an inactive port (**Fig. 2e**) or to ChR2-lacking littermate controls (**Fig. 2f**), indicating excitation of VTA^GG^ projections to LHb can be reinforcing. To test whether release of GABA or of glutamate from VTA^GG^ terminals contributes to this reinforcement, we co-expressed ChR2 in combination with a Cre-dependent Cas9 to selectively delete the gene encoding VGAT in VGLUT2+ neurons, or a Flp-dependent Cas9 vector to selectively delete the gene encoding VGLUT2 in VGAT+ neurons (**Fig. 2g**). Loss of VGAT disrupted GABA release from VTA^GG^ neurons, while loss of VGLUT2 disrupted glutamate release from VTA^GG^ neurons (**Supplemental Fig. 1a-d**). Behaviorally, the selective deletion of glutamate release from VTA^GG^ neurons did not alter optogenetic self-stimulation of VTA^GG^ terminals in LHb, while deletion of GABA release abolished it (**Fig. 2h, Supplemental Data Fig 1e**). This indicates that GABA release, but not glutamate release, is necessary for self-stimulation of VTA^GG^ projections to LHb, consistent with prior works showing LHb inhibition promotes reward^12,26^.

**Figure 2.**
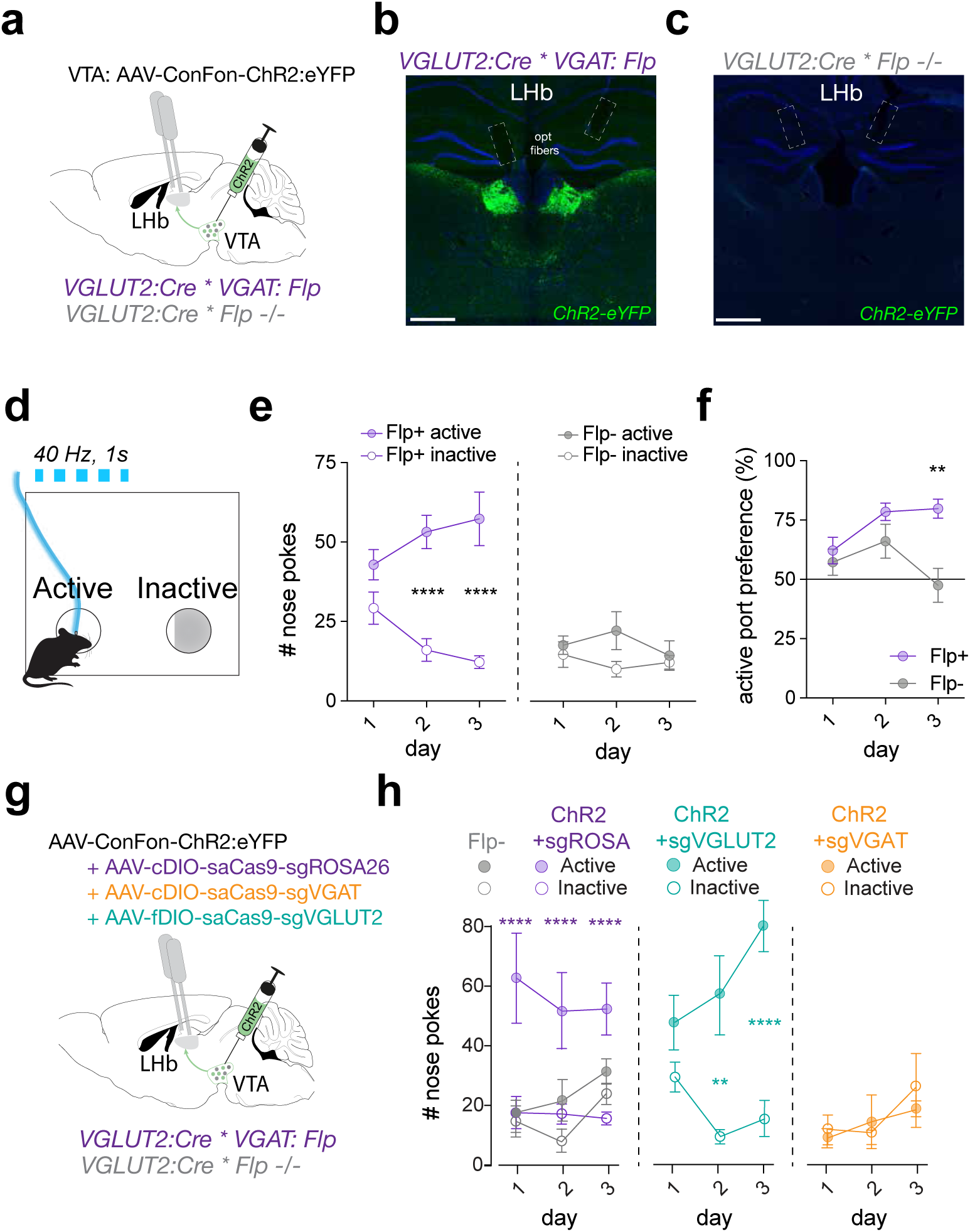
The reinforcing effects of VTA^GG^ input to LHb require GABA release. (A) Strategy to express ChR2:eYFP in VTA^GG^ neurons, plus optic fibers implanted bilaterally above LHb. (B) Terminal expression of ChR2 in LHb of VGLUT2-Cre; VGAT-Flp mice and (C) control mouse lacking Flp; scale = 500 um. (D) Schematic of optogenetic self-stimulation assay. Nose pokes into the active port delivered 40 Hz 1s blue (473 nm) light plus a 1s tone, while nose pokes into the inactive port delivered a 1s tone alone. (E) ChR2-expressing mice (Flp+; n=10) nose poked at greater amounts in the active port compared to the inactive port (Two-Way ANOVA, main effect of port-type: F_1,9_=29, p=0.0004; day x port interaction: F_2,18_=9.3, p=0.002; Bonferroni multiple comparisons between active vs. inactive ports: day 2, t_18_=6.9, p<0.0001; day 3, t_18_=8.4, p<0.0001). Flpo-/-control mice (Flp-; n=9) nose poked at equal amounts across both ports (Two-Way ANOVA, main effect of port-type: F_1,8_=2.7, p=0.14). (F) Flp+ mice showed a higher percent preference for active port than Flp-mice (Two-Way ANOVA, main effect of group: F_1,17_=8.3, p=0.01) that grew over days (day x group interaction: F_2,34_=4.5, p=0.02; Bonferroni multiple comparisons between Flp+ and Flp-preference on day 3: t_13_=3.9, p<0.005). (G) Strategy to express ChR2 and a CRISPR-Cas9 AAV targeted to targeted to the control ROSA26 locus (sgROSA), or to disrupt expression of VGAT (sgVGAT) or VGLUT2 (sgVGAT) in VTA^GG^ neurons; plus optic fibers implanted bilaterally above LHb. (H) Active vs. inactive nose pokes in self-stimulation assay (same as depicted in panel D) for all four groups. Flp-/-control mice (Flp-; n=11) poked equally between ports (Two-Way ANOVA, main effect of port-type: F_1,10_=2.7, p=0.16), sgRosa26 control mice preferred the active port (n=9): Two-Way ANOVA, main effect of port-type: F_1,8_=13, p=0.007; day x port interaction: F_2,16_=0.8, p=0.47; Bonferroni multiple comparisons between active vs. inactive ports: day 1, t_16_=7.3, p<0.0001; day 2, t_16_=5.7, p<0.0001; day 3, t_16_=5.8, p<0.0001); sgVGLUT2 mice preferred the active port (n=9): Two-Way ANOVA, main effect of port-type: F_1,8_=12.4, p=0.008; day x port interaction: F_2,16_=5.7, p=0.01; Bonferroni multiple comparisons between active vs. inactive ports: day 1, t_16_=0.95, p>0.99; day 2, t_16_=3.5, p<0.009; day 3, t_16_=5.7, p<0.0001); sgVGAT mice showed no preference for active port (n=11): Two-Way ANOVA, main effect of port-type: F_1,10_=3.5, p=0.09; day x port interaction: F_2,20_=2.4, p=0.11). Data are shown as mean +/-SEM; *p<0.05, **p<0.01, ***p<0.001, ****p<0.0001.

### Bidirectional, state-dependent effects of GABA-glutamate co-transmission

We next tested how a GPCR-based chemogenetic approach to bidirectionally alter LHb state influenced the net effect of VTA^GG^ input, both *ex vivo* and *in vivo*. We expressed ChR2 in VTA^GG^ neurons plus either excitatory hM3Dq or inhibitory hM4Di receptors in LHb glutamate neurons, allowing us to activate VTA^GG^ inputs in the presence or absence of clozapine-N-oxide (CNO) which activates these receptors (**Fig. 3a-b**). We first observed that, in the absence of optogenetic stimulation, bath application of CNO reliably depolarized hM3Dq-expressing cells and hyperpolarized hM4Di-expressing cells (**Fig. 3c-f**). Next, oPSPs were recorded before and after bath application of CNO. In hM3Dq-expressing cells, oPSPs became smaller during CNO application, whereas oPSPs became more positive in hM4Di-expressing cells, causing the effect of VTA^GG^ stimulation to shift from net inhibitory to net excitatory.

**Figure 3.**
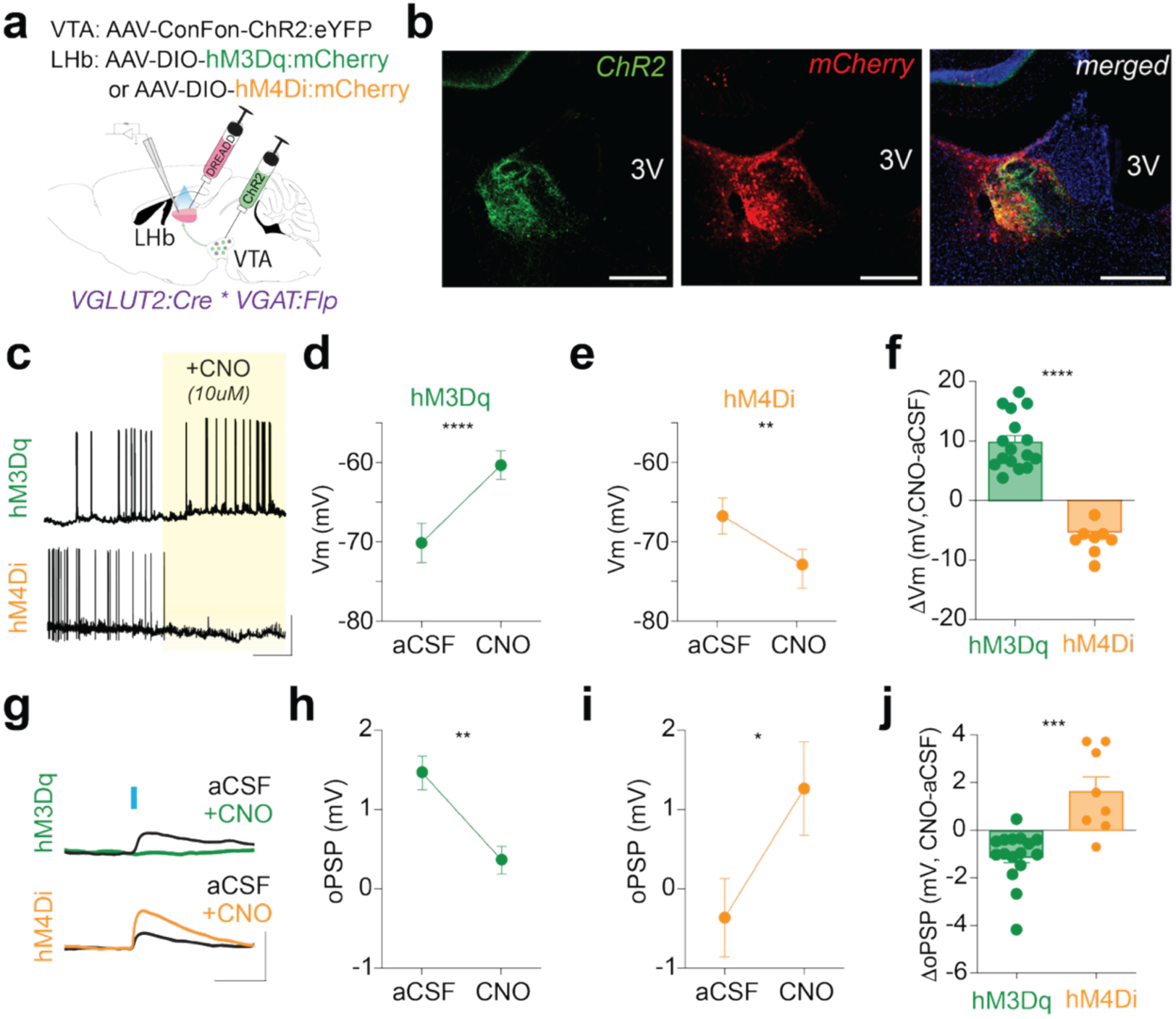
Bidirectional chemogenetic manipulation of LHb reveals state-dependent inhibitory or excitatory effects of VTA^GG^ stimulation ex vivo. (A) Strategy to express ChR2:eYFP in VTA^GG^ neurons, plus designer receptors exclusively activated by designer drug (DREADD) in LHb glutamate neurons. (B) ChR2 expression in VTA^GG^ terminals in LHb (green), hM3Dq:mCherry expression in LHb glutamate neurons, and merged image of both channels; scale = 250um. (C) Example whole-cell (I=0) traces from hM3Dq-and hM4Di-expressing LHb cells before and during CNO bath application; scale: x= 100 s, y=25 mV. (D) Membrane potential increased with CNO compared to pre-CNO (aCSF) period for hM3Dq-expressing cells; paired t-test: t_15_=8.5, p<0.0001. (E) Membrane potential decreased during CNO compared to pre-CNO (aCSF) for hM4Di-expressing cells; paired t-test: t_7_=4.9, p=0.002. (F) Change in membrane potential from aCSF to CNO periods for hM3Dq versus hM4Di-expressing cells; unpaired t-test: t_22_=8.4, p<0.0001. (G) Example optogenetic-evoked postsynaptic potential (oPSP) in response to 5ms light pulse in aCSF (black) or after CNO bath application for hM3Dq cell (green) or hM4Di cell (orange); scale: x= 25 ms, y=5 mV. (H) oPSP decreased during CNO compared to pre-CNO (aCSF) in hM3Dq-expressing neurons; paired t-test: t_15_=3.9, p=0.001. (I) oPSP increased during CNO compared to pre-CNO (aCSF) in hM4Di-expressing neurons; paired t-test: t_7_=2.6 p=0.03. (J) Change in oPSP from aCSF to CNO periods for hM3Dq versus hM4Di-expressing cells; unpaired t-test: t_22_=4.7, p=0.0001.

To assess how bidirectional modulation of LHb state altered the net effect of VTA^GG^ on population-level activity *in vivo* we used a similar approach, with the addition of fiber photometry. To validate chemogenetic effects *in vivo* we expressed either hM3Dq or hM4Di in LHb glutamate neurons, treated mice with CNO, and measured Fos induction. CNO increased Fos expression in LHb neurons from mice expressing hM3Dq compared to mCherry, while hM4Di led to a small reduction that was not statistically significant compared to mCherry, perhaps due to floor effects (**Supplemental data Fig 3a-c**).

We next expressed a red-shifted Channelrhodopsin (ChRmine) in VTA^GG^ neurons. And in LHb glutamate neurons we co-expressed GCaMP to record calcium activity with either control (mCherry, **Supplemental Data Fig. 2**), hM3Dq, or hM4Di (**Fig. 4a-d**). Prior to treating mice with CNO, we first measured how activation of VTA^GG^ projections at variable durations alters calcium activity in LHb at baseline. We found that VTA^GG^ stimulation evoked transient decreases in LHb activity, with a slower sustained inhibitory process apparent when stimulation duration was prolonged (**Fig. 4e-h**). We also observed a strong post-stimulation increase in activity, presumably an inhibition-induced rebound effect, as has been previously observed in LHb^12,27–29^. In separate sessions, we tested the effects of VTA^GG^ stimulation before and after CNO treatment in order to assess the effects of bidirectional manipulation of LHb (**Fig. 4i**).

**Figure 4.**
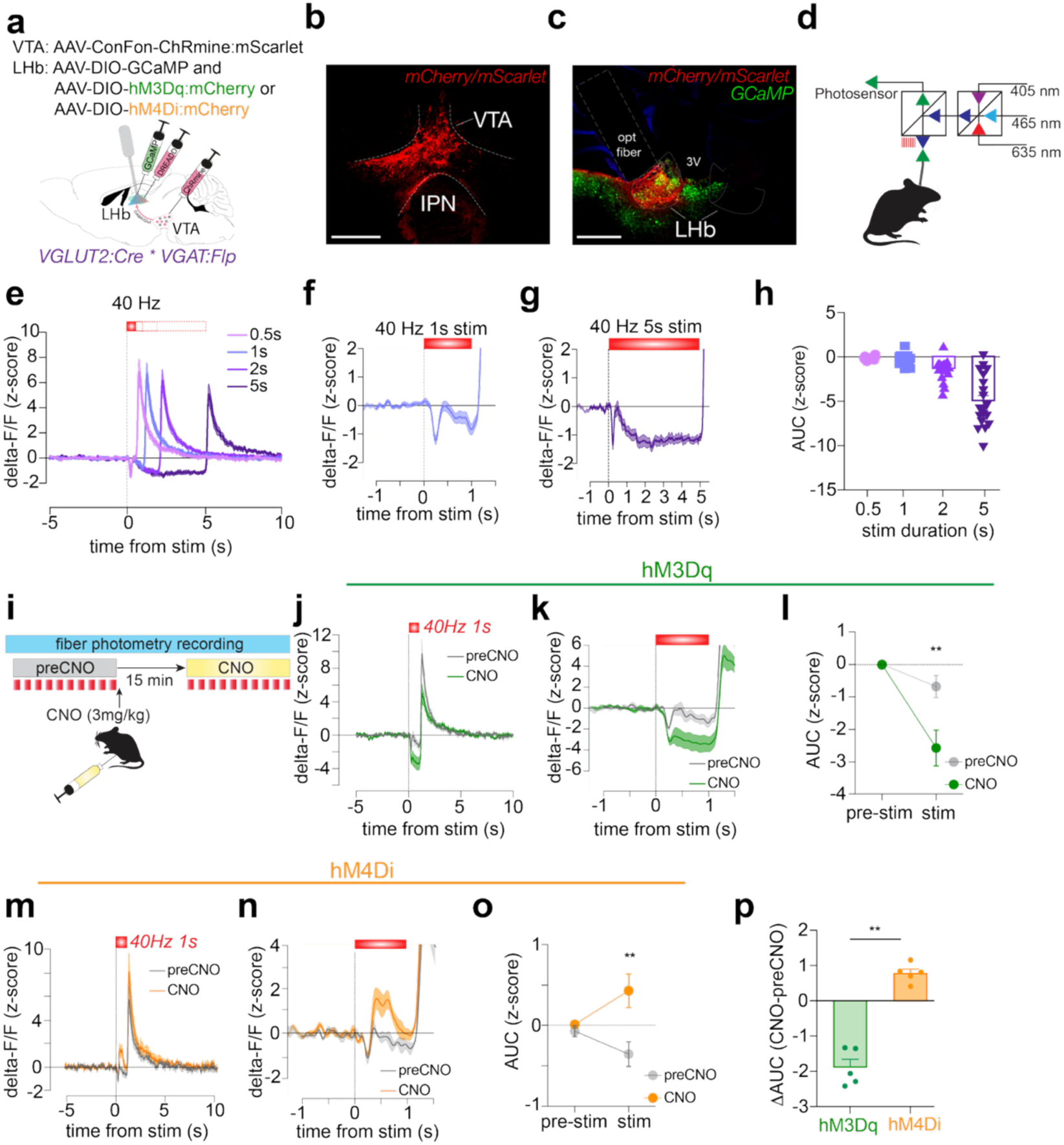
Bidirectional chemogenetic manipulation of LHb reveals state-dependent inhibitory or excitatory effects of VTA^GG^ stimulation in vivo. (A) Strategy to express the red-shifted channelrhodopsin ChRmine:mScarlet in VTA^GG^ neurons, plus co-express GCaMP and DREADD:mCherry (hM3Dq or hM4Di) in LHb glutamate neurons, plus implant an optic fiber above LHb. (B) ChRmine expression in VTA; scale = 500um. (C) Expression of GCaMP and DREADD:mCherry in LHb neurons, along with expression of ChRmine:mScarlet in VTA^GG^ terminals; scale = 500 um. (D) Schematic showing wavelengths used to monitor GCaMP fluorescence by fiber photometry in freely moving mice (465 nm: GCaMP, 405 nm: isosbestic control, 635 nm: optogenetic excitation of VTA^GG^ terminals in LHb). (E) Average delta F/F z-score of GCaMP signals from all mice during optogenetic stimulation (40 Hz) at varying durations reveals stimulus-evoked inhibition followed by post-stimulus rebound activation; (F) zoomed into stimulation period delivered for 1s or (G) 5s. (H) Average area-under-curve (AUC) during stimulation period for the varying stimulation durations; one-way ANOVA: F_3,72_=37.9, p<0.0001. (I) Schematic showing clozapine-N-oxide (CNO) sessions: 10 trials of optogenetic stimulation were delivered during baseline (preCNO), and 10 trials of the same stimulation delivered after i.p. CNO injection. (J) Average z-score of hM3Dq mice (n=5) during 40Hz, 1s stimulation; (K) zoomed in stimulation period for preCNO (gray) and CNO (green) trials. (L) hM3Dq activation led to increased stim-evoked inhibition; two-way ANOVA; main effect of stim period: F_1,8_=21.9, p=0.002; main effect of CNO: F_1,8_=8.7, p=0.02; interaction of stim period x CNO: F_1,8_=7.3, p=0.03; Holm-Sidak multiple comparisons test, preCNO vs. CNO during pre-stim: t_16_=0.02, p=0.98; Holm-Sidak multiple comparisons test, preCNO vs. CNO during stim: t_16_=3.98, p=0.002. (M) Average z-score of hM4Di mice (n=5) during 40Hz, 1s stimulation; (N) zoomed in stimulation period for pre-CNO (gray) and CNO (orange) trials. (O) hM4Di activation led to decreased stim-evoked inhibition and revealed stim-evoked excitation; two-Way ANOVA, main effect of stim period: F_1,8_=0.25, p=0.63; main effect of CNO: F_1,8_=10.2, p=0.01; interaction of stim period x CNO: F_1,8_=6.8, p=0.03; Holm-Sidak multiple comparisons test, preCNO vs. CNO during pre-stim: t_16_=0.43, p=0.67; Holm-Sidak multiple comparisons test, preCNO vs. CNO during stim: t_16_=4.1, p=0.002. (P) Change in AUC after CNO compared to preCNO trials for hM3Dq compared to hM4Di mice; Mann-Whitney test, U=0, p=0.008. Data shown as mean +/-SEM; **p<0.01.

Activation of hM3Dq in LHb neurons caused the effects of VTA^GG^ stimulation to become even more inhibitory (**Fig. 4j-l**). Conversely, activation of hM4Di in LHb neurons caused VTA^GG^ input to become either less inhibitory or flip to excitatory (**Fig. 4m-o**). These results indicate that VTA GABA-glutamate co-transmission can flexibly shift signs between inhibition and excitation in a manner contingent on the activity state of the postsynaptic neuron (**Fig. 4p**).

### State-dependent reinforcing effect of VTA^GG^ stimulation

We show above that the reinforcing effect of VTA^GG^ reinforcement is dependent on inhibitory GABA transmission onto LHb, and that VTA^GG^ stimulation tends to inhibit LHb more strongly when LHb was depolarized, whereas inhibition reversed to excitation when LHb was hyperpolarized. We reasoned that the reinforcing effect of VTA^GG^ terminals would likewise change as a function of LHb activity state. To test this, we used a similar chemogenetic strategy to bidirectionally manipulate LHb while mice were given the opportunity to self-stimulate VTA^GG^ projections to LHb (**Fig. 5a-b, Supplemental data Fig. 3f**). All ChR2-expressing groups reliably self-stimulated VTA^GG^ inputs during the first 3 days (vehicle) compared to littermates that lacked ChR2 expression (**Fig. 5c-g**). On days 4-6, mice were treated with CNO prior to self-stimulation sessions to activate either hM3Dq or hM4Di.

**Figure 5.**
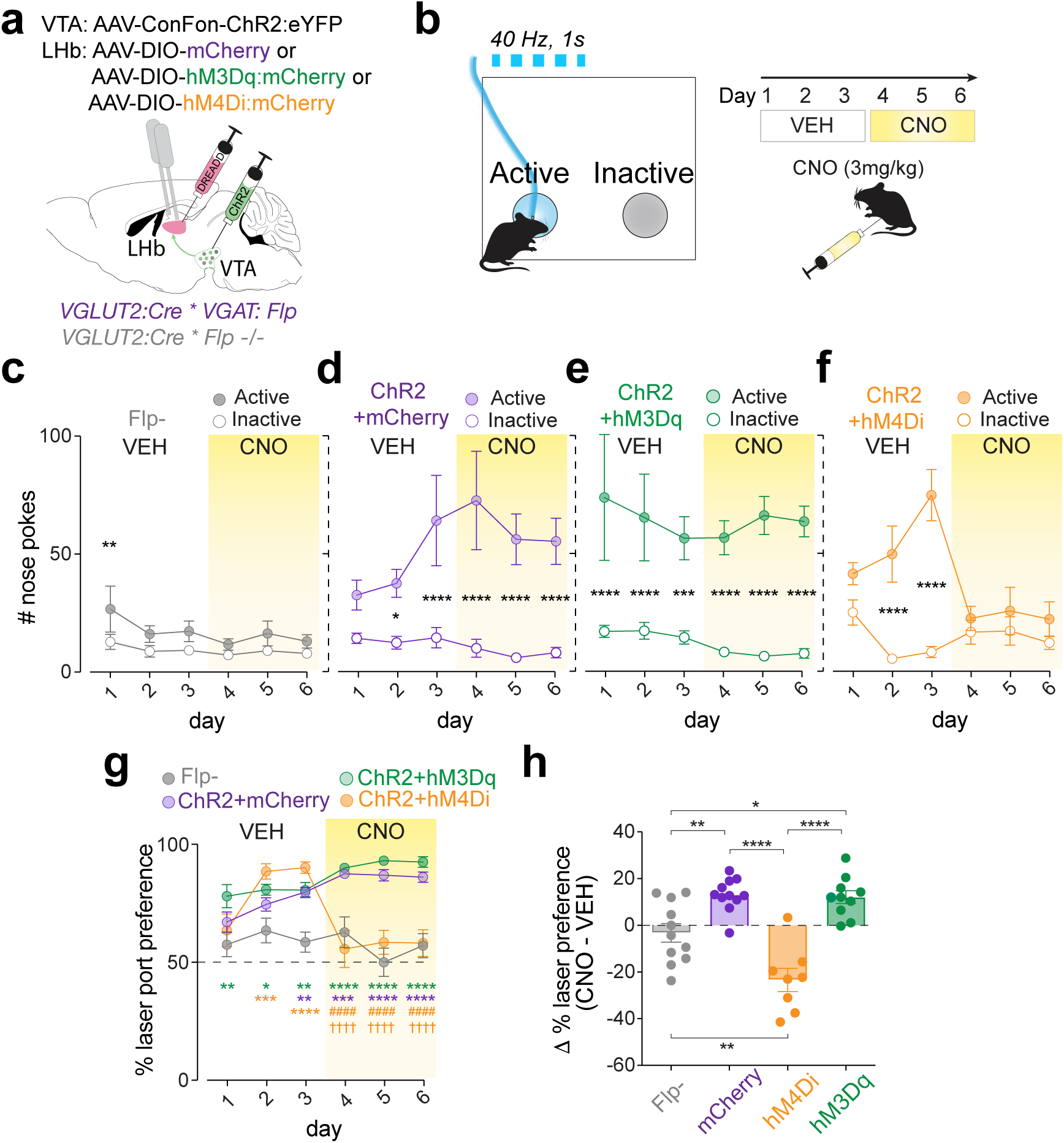
Bidirectional chemogenetic manipulation of LHb reveals state-dependent reinforcing effects of VTA^GG^ projections. (A) Strategy to express ChR2:eYFP in VTA^GG^ neurons, plus DREADD:mCherry (hM3Dq or hM4Di) or mCherry control in LHb, plus optic fibers implanted bilaterally above LHb. (B) Schematic of optogenetic self-stimulation assay. Vehicle (0.5% DMSO) was injected (i.p.) 15 min prior to sessions on days 1-3 and CNO was injected (i.p.) 15 min prior to sessions on days 4-6. (C) Flp-negative control mice that do not express ChR2 (n=11) poked equally between ports (two-way ANOVA, main effect of port-type: F_1,10_=8.7, p=0.01; main effect of day: F_5,50_=1.5, p=0.19; day x port interaction: F_5,50_=0.7, p=0.63; Bonferroni multiple comparisons between active vs. inactive ports: day 1, t_50_=3.5, p=0.01). (D) ChR2-and mCherry (DREADD control)-expressing mice (ChR2+mCherry; n=11) nose poked in greater amounts at the active port compared to the inactive port (two-way ANOVA, main effect of port-type: F_1,10_=30, p=0.0002; day x port interaction: F_5,50_=3.4, p=0.009; Bonferroni multiple comparisons between active vs. inactive ports: day 2, p=0.04; days 3-6, p<0.0001). (E) ChR2-and hM3Dq-expressing mice (ChR2+hM3Dq; n=10) nose poked at greater amounts in the active port compared to the inactive port (two-way ANOVA, main effect of port-type: F_1,9_=26, p=0.0006; day x port interaction: F_5,45_=0.4, p=0.79; Bonferroni multiple comparisons between active vs. inactive ports: days 1-2, p<0.0001; day 3, p=0.0005; days 4-6, p<0.0001). (F) ChR2-and hM4Di-expressing mice (ChR2+hM4Di; n=8) nose poked at greater amounts in the active port compared to the inactive port, only on VEH days (two-way ANOVA, main effect of port-type: F_1,7_=59, p=0.0001; main effect of day: F_5,35_=3.4, p=0.01; day x port interaction: F_5,35_=8.9, p<0.0001; Bonferroni multiple comparisons between active vs. inactive ports: days 2-3, p<0.0001). (G) Percent preference for the active, laser-paired port (two-way ANOVA, main effect of group: F_3,36_=33, p<0.0001; main effect of day: F_5,180_=3.8, p=0.003; day x group interaction: F_15,180_=6.2, p<0.0001; Bonferroni multiple comparisons between each group: day 1: hM3Dq vs. Flp-, p<0.01; day 2: hM3Dq vs. Flp-, p<0.05; hM4Di vs. Flp-, p<0.001; day 3: hM3Dq vs. Flp-, p<0.01; hM4Di vs. Flp-, p<0.0001; ChR2+mCherry vs. Flp-, p<0.01; day 4: hM3Dq vs. Flp-, p<0.0001; ChR2+mCherry vs. Flp-, p<0.001; ChR2+mCherry vs. hM4Di, p<0.0001; hM3Dq vs. hM4Di, p<0.0001). day 5: hM3Dq vs. Flp-, p<0.0001; ChR2+mCherry vs. Flp-, p<0.0001; ChR2+mCherry vs. hM4Di, p<0.0001; hM3Dq vs. hM4Di, p<0.0001; day 6: hM3Dq vs. Flp-, p<0.0001; ChR2+mCherry vs. Flp-, p<0.0001; ChR2+mCherry vs. hM4Di, p<0.0001; gM3Dq vs. hM4Di, p<0.0001). * vs. Flp-, ^#^ vs. ChR2+mcCerry, ^†^vs. hM3Dq. (H) Change in percent preference between CNO and VEH sessions (one-way ANOVA, F_3,36_=33, p<0.0001; Bonferroni multiple comparisons between each group: Flp-vs. ChR2+mCherry, p=0.007; Flp-vs. hM4Di, p=0.002; Flp-vs. hM3Dq, p=0.02; ChR2+mCherry vs. hM4Di, p<0.0001; hM4Di vs. hM3Dq, p<0.0001). Data are shown as mean +/-SEM; *p<0.05, **p<0.01 ***p<0.001, ****p<0.0001;

ChR2-expressing mice with hM3Dq in LHb maintained their rate of self-stimulation (**Fig. 5e**) and showed an increase in preference for the active port on CNO days compared to vehicle days (**Fig. 5h**). However, this did not statistically differ from the mCherry control group using multiple comparisons, perhaps due to a ceiling effect (hM3Dq mice reached ∼95% preference for the laser port by day six). By contrast, self-stimulation was almost abolished by CNO treatment in hM4Di-expressing mice (**Fig. 5f**). Lack of self-stimulation was not due to generalized locomotor deficits, as CNO did not affect movement of hM4Di-expressing mice in an open-field assay, though CNO did reduce movement in hM3Dq-expressing mice in this assay (**Supplemental Data Fig. 3d-e**). Overall, these results suggest that the reinforcing effects of VTA^GG^ stimulation are state-dependent, becoming less reinforcing when LHb activity is suppressed and the effects of VTA^GG^ co-release are shifted toward excitatory.

## Discussion

A majority of individual neurons projecting to the LHb co-transmit inhibitory GABA and excitatory glutamate, including a substantial input from VTA^4,21^. Cotransmission of GABA and glutamate throughout the brain may serve a variety of functions such as promoting neural development, or as a compensatory response to prevent seizures^2,23,30^. However, its function within the brain’s reward circuitry has remained unclear. Here we show that GABA-glutamate projections to LHb are net inhibitory at baseline and can promote reinforcement through release of GABA. However, shifting the activity state of LHb using chemogenetics caused the effect of GABA-glutamate cotransmission to change signs: making its net effect more excitatory in the case of LHb inhibition, and more inhibitory when LHb was excited. We observed this dynamic shift not only in individual postsynaptic cells *ex vivo*, but *in vivo* at a population level using calcium sensors. The state-dependent switch from net inhibition to net excitation abolished the reinforcing effect of VTA^GG^ stimulation. These results demonstrate that the net effects of VTA^GG^ on LHb can dynamically switch signs, and help reconcile conflicting reports of VTA projections to LHb being rewarding in some contexts, but aversive in others^9,31^.

In LHb, shifts from hypoactivity to hyperactivity (or vice versa) are associated with development of neuropsychiatric disorders such as depression, bipolar, compulsions, and addiction^17,32–34^.

Indeed, dysregulation of GABA-glutamate cotransmission in the LHb as a result of early life stress or chronic drug exposure has been associated with dysregulated mood and motivation^35–37^. Given that inhibitory interneurons within LHb are sparse^25^, GABA-glutamate cotransmission may represent an alternative strategy to homeostatically prevent runaway hyperactivity or hypoactivity, protecting against the development of these disorders.

Prior work showed that chronic manipulations (days to weeks) induce presynaptic changes that modify the ratio of GABA and glutamate corelease onto LHb neurons^36,38^. Our results complement those studies and indicate that rapid (sub-second to minutes) postsynaptic changes determine how LHb neurons respond to coreleased GABA and glutamate. We propose a homeostatic model whereby the instantaneous activity state of the postsynaptic cell determines whether VTA^GG^ input is net excitatory or inhibitory. This follows from the principle that the driving force acting on a permeant ion is greater when the neuronal membrane potential is further from that ions reversal potential. Thus, in relatively more depolarized LHb neurons, GABA receptor activation will result in larger hyperpolarizing IPSPs, while glutamate receptor activation will trigger smaller EPSPs. Conversely, in more hyperpolarized LHb neurons, GABA-mediated IPSPs will be relatively smaller (less hyperpolarizing) and glutamate-mediated EPSPs larger. This model is consistent with prior observations using single unit recordings from LHb neurons, finding that aversive footshock tended to activate LHb neurons that were quiescent or firing slowly, but inhibit LHb neurons that were firing faster^27^.

While this model explains our basic physiological and behavioral findings, the effects of GABA-glutamate co-transmission in this circuit are undoubtedly much richer. The passive integration of synchronized EPSPs and IPSPs that occur following glutamate and GABA corelease will depend on the dynamic number and kinetics of receptors present on the membrane and their relative locations along somatodendritic compartments. These properties will also influence their ability to engage non-linear processes such as the recruitment of voltage-dependent cation channels or shunting inhibition^39–42^. In other circuits GABA-glutamate cotransmission has led to divergent patterns of short-term depression that allow for frequency-dependent filtering^43^. Moreover, VTA^GG^ transmission in LHb may act not only on postsynaptic ionotropic receptors but also on GPCRs and on presynaptic sites.

We used chemogenetic manipulations to bidirectionally alter postsynaptic LHb activity states to test how activity state alters the effects of GABA-glutamate co-transmission on physiology and behavior. We found that changes in postsynaptic activity state could flip the sign of GABA-glutamate co-transmission between excitation and inhibition. Future studies could test this hypothesis over longer timescales, for example using chronic stress, pain, or psychoactive drugs to induce adaptations in LHb activity state^35–37^. Future works may also consider shorter timescales. For example, how transient changes in LHb activity due to changing expectations for positive or negative outcomes influence the effects of VTA^GG^ transmission on postsynaptic neurons and behavior. Indeed, a recent pre-print shows that pairing optogenetic stimulation of GABA-glutamate neurons in the entopeduncular nucleus with a rewarding or aversive stimulus could change the effects of these stimuli on *in vivo* dopamine release, and in a manner that correlated with subsequent *ex vivo* effects on the sign of evoked GABA-glutamate cotransmission in LHb^44^.

Overall, our results demonstrate that GABA-glutamate cotransmission can drive net excitatory or inhibitory responses that depend on the state of the postsynaptic cell. This represents a novel homeostatic mechanism by which brain reward circuitry may be regulated to stabilize reward-seeking behaviors, motivation, or mood.

## Supplemental Figures

**Supplemental Figure 1.**
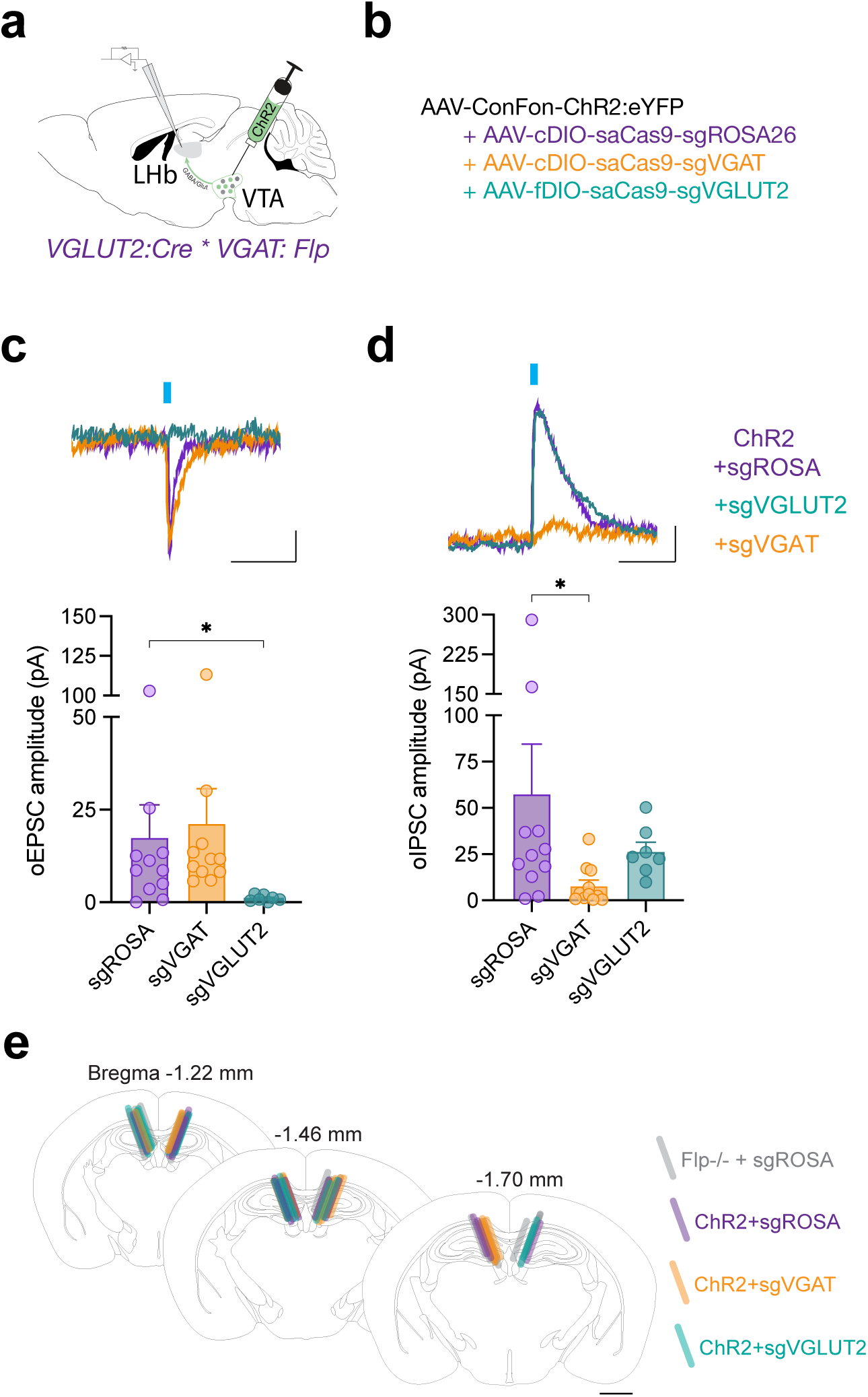
to accompany. **Figure 2. CRISPR/Cas9 deletion of VGAT or VGLUT2, respectively, disrupts evoked GABA or evoked glutamate release from VTA^GG^ projections to LHb ex vivo.** (A) Strategy to express channelrhodopsin (ChR2) and (B) a CRISPR-Cas9 AAV targeted to the ROSA26 control locus (sgROSA), or to disrupt expression of VGAT (sgVGAT) or VGLUT2 (sgVGAT) in VTA^GG^ neurons. (C) Optogenetic-evoked excitatory postsynaptic currents (oEPSCs) in all groups (Kruskal-Wallis test: KW=12.6, p=0.002; Dunn’s multiple comparisons test between sgROSA and sgVGAT mice: p=0.65, and between sgROSA and sgVGLUT2 mice: p<0.02); scale: x=50 ms, y=2.5 pA. (D) Optogenetic-evoked inhibitory postsynaptic currents (oIPSCs) in all groups (Kruskal-Wallis test: KW=9.6, p=0.008; Dunn’s multiple comparisons test between sgROSA and sgVGAT mice: p=0.01, and between sgROSA and sgVGLUT2 mice: p>0.99); scale: x=50 ms, y=5 pA. (E) Map of optic fiber placements in mice used in the 2-nose-poke assay shown in Figure 2G-H; scale bar = 1mm. Data shown as mean +/-SEM or as individual points; *p<0.05, **p<0.01.

**Supplemental Figure 2.**
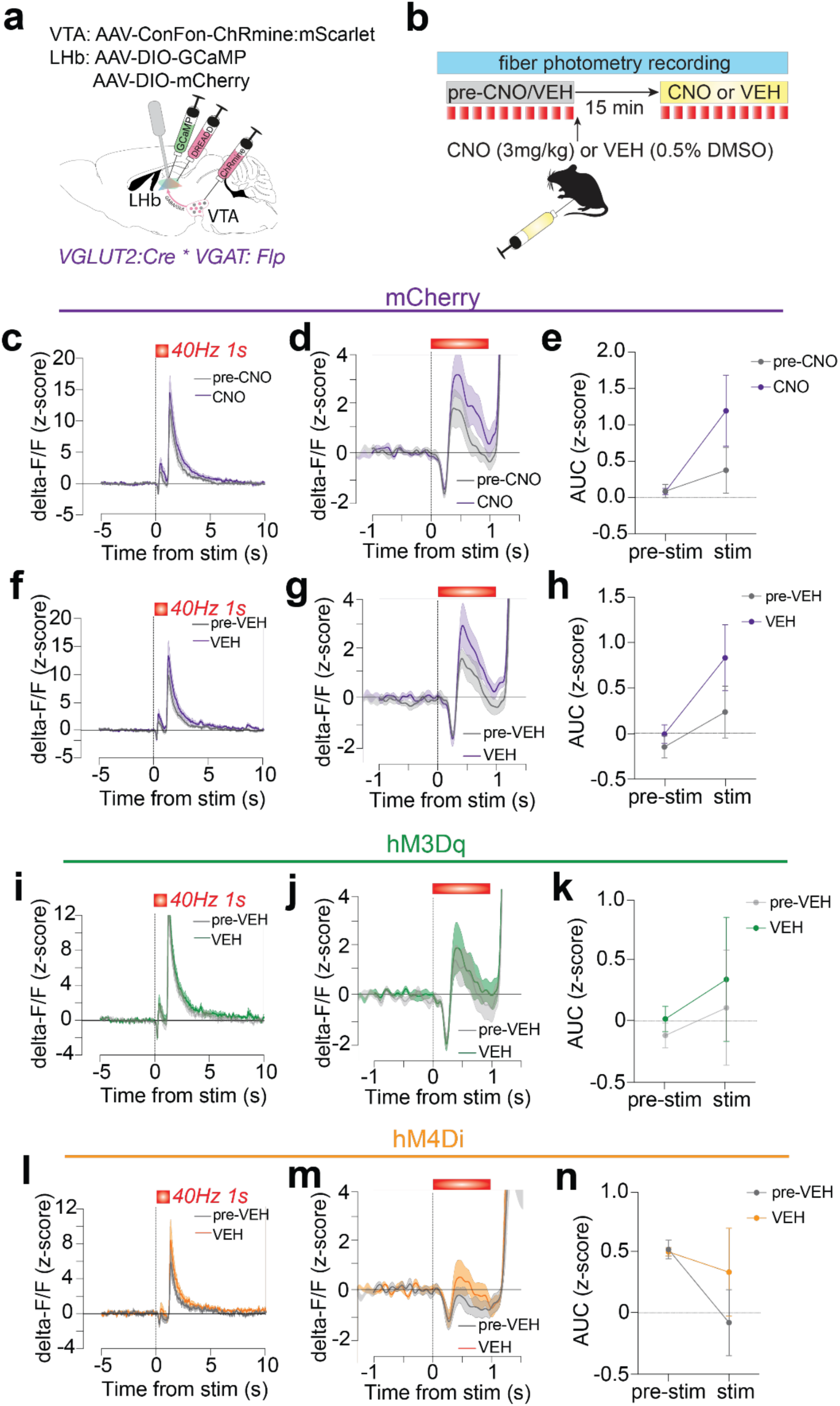
to accompany. **Figure 4. Control groups show no significant effects of CNO or VEH on LHb activity evoked by VTA^GG^ neurons in vivo.** (A) Strategy to express red-shifted channelrhodopsin ChRmine:mScarlet in VTA^GG^ neurons, plus co-express GCaMP and DREADD:mCherry (hM3Dq, hM4Di, or mCherry control) in LHb glutamate neurons, plus an optic fiber implanted above LHb. (B) Schematic showing clozapine-N-oxide (CNO) or vehicle (VEH) sessions: 10 trials of optogenetic stimulation were delivered during baseline (pre-CNO, pre-VEH), and 10 trials of the same optogenetic stimulation were delivered after systemic i.p. CNO or VEH injection. (C) Average delta-F/F z-score of mCherry mice (n=6) during 40Hz, 1s stimulation, (D) zoomed in to during stimulation period, for pre-CNO (gray) and CNO (purple) trials. (E) Average AUC during pre-stimulation (pre-stim) and stimulation periods (stim) for mcherry mice for pre-CNO and CNO trials. Two-Way ANOVA, Main effect of stim period: F_1,15_=13.8, p=0.002. Main effect of CNO: F_2,15_=1.3, p=0.3. Interaction of stim period x CNO: F_2,15_=1.9, p=0.19. (F) Average z-score of mCherry mice during 40Hz, 1s stimulation, during VEH session (G) zoomed in to during stimulation period, for pre-VEH (gray) and VEH (purple) trials. (H) Average AUC during pre-stimulation (pre-stim) and stimulation periods (stim) for mcherry mice for pre-VEH and VEH trials. Two-Way ANOVA, Main effect of stim period: F_1,15_=10, p=0.006. Main effect of VEH: F_2,15_=.8, p=0.46. Interaction of stim period x CNO: F_2,15_=0.5, p=0.6. (I) Average z-score of hM3Dq mice (n=5) during 40Hz, 1s stimulation, during VEH session (J) zoomed in to during stimulation period, for pre-VEH (gray) and VEH (green) trials. (K) Average AUC during pre-stimulation (pre-stim) and stimulation periods (stim) for HM3Dq mice for pre-VEH and VEH trials. Two-Way ANOVA, Main effect of stim period: F_1,4_=0.3, p=0.59. Main effect of VEH: F_1,4_=.7, p=0.45. Interaction of stim period x CNO: F_1,4_=0.8, p=0.43. (L) Average z-score of hM4Di mice (n=5) during 40Hz, 1s stimulation, during VEH session (M) zoomed in to during stimulation period, for pre-VEH (gray) and VEH (orange) trials. (N) Average AUC during pre-stimulation (pre-stim) and stimulation periods (stim) for hM4Di mice for pre-VEH and VEH trials. Two-Way ANOVA, Main effect of stim period: F_1,4_=59, p=0.001. Main effect of VEH: F_1,4_=1.8, p=0.25. Interaction of stim period x CNO: F_1,4_=4, p=0.12. Data are shown as mean +/-SEM.

**Supplemental Figure 3.**
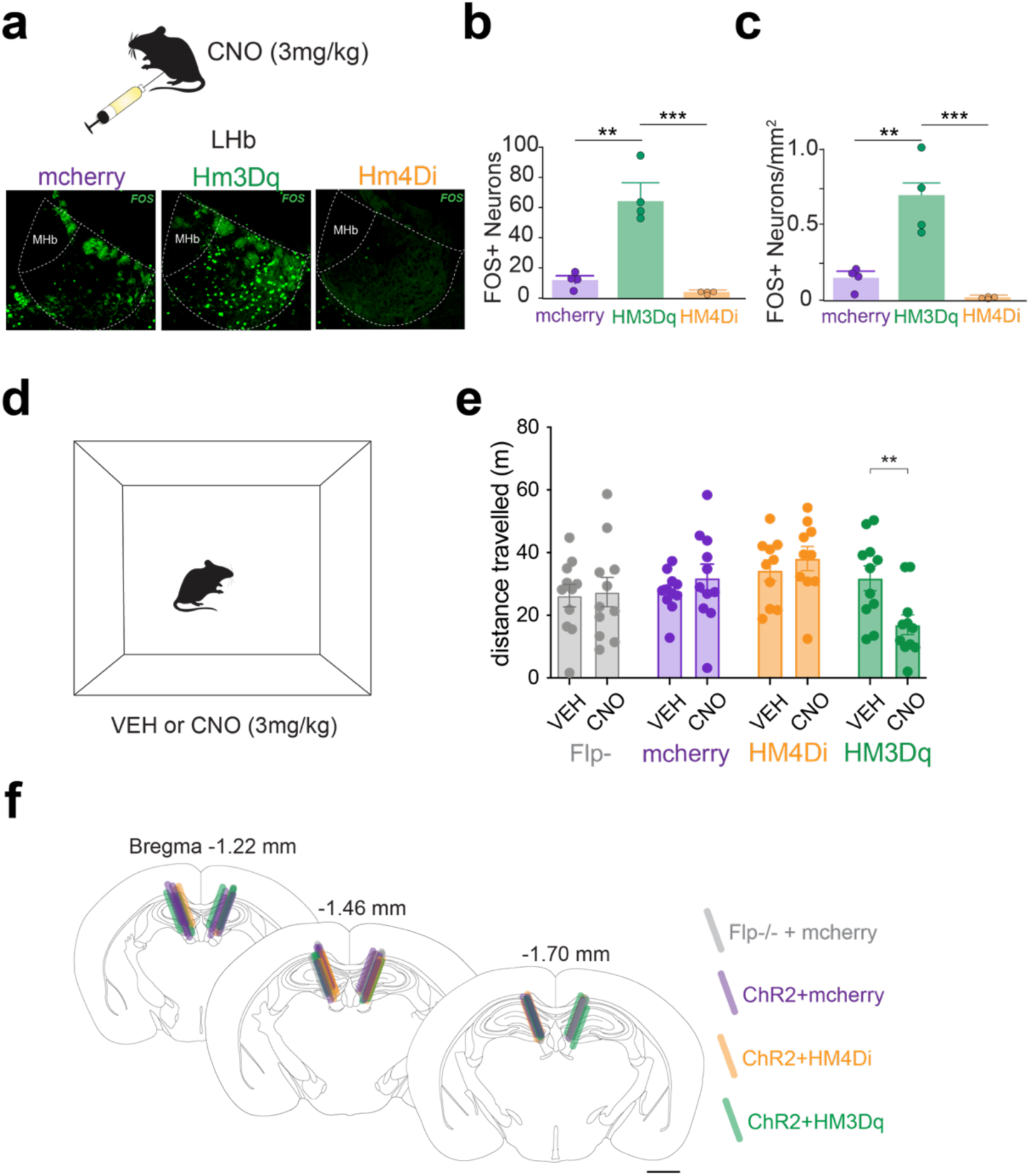
to accompany. **Figure 5. Effects of bidirectional chemogenetic manipulation of LHb on Fos expression and in an open-field test.** (A) Fos expression in LHb after CNO injection (3mg/kg, i.p.) in mice expressing mCherry (control), hM3Dq:mCherry, or hM4Di:mCherry mice. (B) Total Fos+ neurons in LHb; one-way ANOVA, F_2,9_=34, p<0.0001; Bonferroni multiple comparisons between each group: mCherry vs. hM3Dq, p=0.0004; mCherry vs. hM4Di, p=0.62; hM4Di vs. hM3Dq, p<0.0001). (C) Fos+ neuron density in LHb; one-way ANOVA, F_2,9_=23, p=0.0003; Bonferroni multiple comparisons between each group: mCherry vs. hM3Dq, p=0.002; mCherry vs. hM4Di, p=0.48; hM4Di vs. hM3Dq, p<0.0003). (D) Open-field locomotor activity during a 20 min session after vehicle (VEH) or CNO (3 mg/kg, i.p.) injection. (E) Distance travelled during open-field sessions; two-way ANOVA: main effect of drug: F_1,39_=0.5, p=0.48; main effect of DREADD expression: F_3,39_=2.5, p=0.06; drug x expression group interaction: F_3,39_=5.3, p=0.004; Bonferroni multiple comparisons between VEH and CNO sessions: hM3Dq group, p=0.002. (F) Map of optic fiber placements in 2-nose-poke assay shown in Figure 5; scale bar = 1mm. Data shown as mean +/-SEM; *p<0.05, **p<0.01 ***p<0.001.

## Methods

### EXPERIMENTAL SUBJECTS

Male and female mice were bred at University of California San Diego (UCSD) and group-housed on a 12-hour light/dark cycle. Food and water were available *ad libitum* unless otherwise noted. *Slc17a6^+/Cre^* (VGLUT2-IRES-cre) mice and *Slc32a1^Flp+/-^* (VGAT-Flp) mice were initially obtained from Jackson Laboratory (Stocks: 018147 and 029591, respectively), and were crossed to generate a dual transgenic, VGLUT2-cre/VGAT-Flp line. All experiments were performed on mice at least 6 weeks of age and in accordance with protocols approved by the UCSD Institutional Animal Care and Use Committee.

### METHOD DETAILS

#### Viral Production

Production of AAV1-ConFon-ChR2-EYFP, AAV1-FLEX-EYFP, AAV1-FLEX-SaCas9-Ug-sgVGAT, AAV1-FRT-SaCas9-U6-sgVGLUT2, and AAV1-FLEX-SaCas9-U6-sgRosa26 were as previously described^1^. pAAV shuttle plasmids were co-transfected with packaging plasmid pDG1^2^ into HEK293T/17 cells (ATCC) and viral particles were purified by cesium chloride gradient centrifugation. Viral particles were resuspended in Hank’s balanced saline solution and titers were calculated using gel electrophoresis and densitometry against a known standard.

#### Stereotactic Surgeries

Mice >6 weeks old were anesthetized with isoflurane (4% for induction; 1-2% for maintenance) and placed in a stereotaxic frame (Kopf Instruments). 300 nL total volume of AAV8-ConFon-ChR2-eYFP (2.5 x 10^13^ gc/mL; Addgene 55645) was injected into the VTA of VGLUT2-Cre/VGAT-Flp or VGLUT2-Cre/VGAT-Flp^-/-^ mice (Distance from Bregma in mm:-3.4 AP; +0.35 ML;-4.4 DV) at 100 nl/min using a glass pipette attached to a microinjector (Nanoject 3, Drummond Scientific). Following viral infusion, the injector tip was kept in place for 10 min before slowly retracting. For CRISPR/Cas9 experiments, AAV vectors were made in-house (Zweifel lab) as described above, and AAV1-ConFon-ChR2-EYFP (8 x 10^12^ gc/mL) was combined with either AAV1-FLEX-SaCas9-U6-sgVGAT (1.2 x 10^12^ gc/mL) AAV1-FRT-SaCas9-U6-sgVGLUT2 (1.8 x 10^12^ gc/mL), or AAV1-FLEX-SaCas9-U6-sgRosa26 (1.5 x 10^12^ gc/mL) such that ChR2 constituted 1/4^th^ of the total volume and the respective SaCas9 constituted 3/4^th^ of the total volume (400 nL volume). For DREADD experiments, AAVDJ-EF1⍺-DIO-hM3Dq-mCherry (6.56 x 10^11^ gc/mL; Salk Institute), AAVDJ-EF1⍺-DIO-hM4Di-mCherry (6.04x 10^11^ gc/mL; Salk Institute), or AAVDJ-EF1⍺-DIO-mCherry (2 x 10^12^ gc/mL; UNC) were also injected into LHb during the same surgery (150nl volume; Distance from Bregma in mm:-1.46 AP; +/-1.4 ML;-2.4 DV; 20° medio-lateral angle). For optogenetic self-stimulation experiments, optic fibers (200µm core; RWD) were subsequently placed bilaterally above LHb, 0.2 mm dorsal to the injection site, and optic fibers were secured with 2-4 skull screws and dental cement (Lang Dental Mfg.).

For fiber photometry experiments, AAV8-nEF-Con/Fon-ChRmine-oScarlet (1.9 x 10^13^ gc/mL; Addgene 137159) was injected into VTA as above to express a red-shifted ChR2 in VTA^GG^ neurons. AAV5-syn-FLEX-jGCaMP8f-WPRE (1.97 x 10^13^ gc/mL; Addgene 162379) was combined with either hM3Dq, hM4Di, or mCherry DREADDs (same as above) such that each virus constituted 1/2^th^ of the total volume (200nl), and then injected into LHb at the same coordinates as above. An optic fiber (400µm core; RWD) was subsequently placed above LHb, 0.05 mm dorsal to the injection site and then secured with 2-4 skull screws and dental cement (Lang Dental Mfg.).

For all surgeries, mice were treated with topical antibiotic and carprofen (5mg/kg; s.c.; Rimadyl) immediately following surgery and 24 hr later. Mice were allowed 3-5 weeks for recovery before experiments began.

#### Immunohistochemistry

Mice were deeply anesthetized with sodium pentobarbitol (200mg/kg; i.p.; VetOne) and transcardially perfused with 40-50 mL of phosphate-buffered saline (PBS) followed by 6-70 mL of 4% paraformaldehyde (PFA) at ∼7 mL/min. Brains were extracted and stored in 4% PFA overnight followed by cryoprotection in 30% sucrose PBS for 48-72 hr at 4° C. Brains were subsequently flash frozen in isopentane and dry ice, and stored at-80° C. 40µm sections were cut using a cryostat (Leica) and stored in 0.01% sodium azide PBS. For immunostaining, sections were blocked with 4% normal donkey serum (Jackson ImmunoResearch) in PBS containing 0.2% Triton-X 100 for 1-2 hrs at room temperature. Sections were then incubated in rabbit anti-GFP (1:2000; Invitrogen A11122), or chicken anti-GFP (1:2000; A10262) and rabbit anti-dsRed (1:2000; Clontech 632496) to stain for ChRmine-oScarlet or for DREADD-mCherry overnight at 4° C. Following primary incubation, sections were rinsed in PBS 3 times for 10 min each and subsequently incubated in secondary antibodies conjugated to Alexa 488 (Donkey anti-rabbit; 1:400) or Alexa 488 (Donkey anti-chicken;1:400) and Alexa 594 (Donkey anti-rabbit; 1:400) (Jackson ImmunoResearch) for 2 hr at room temperature. Sections were then rinsed in PBS 3 times for 10 min each, mounted onto glass slides, and coverslipped with Fluoromount-G mounting medium (Southern Biotech) containing 0.5µg/mL DAPI (Sigma). For CNO-mediated c-Fos experiments, rabbit anti-cFos (1:2000; Cell Signaling 2250S) was used followed by a secondary antibody conjugated to Alexa 488 (Donkey anti-rabbit, 1:400).

Images were taken using a widefield epifluorescent microscope (Zeiss AxioObserver).

Tiled images were taken at 10x magnifications using appropriate filters (TXRED, FITC, DAPI), and identical acquisition settings across all slides. Optic fiber placements were mapped onto corresponding coronal sections in the Paxinos Mouse Brain Atlas using Adobe Illustrator. Mice were excluded from experiments if there was substantial spread of ChR2 outside of VTA (beyond 10% volume, n=1), DREADD or GCaMP outside of LHb (n=5) or if optic fibers were placed outside of LHb, either ventrally, or medio-laterally (n=4).

#### Electrophysiological recordings in mouse brain slices

Adult mice 10-14 weeks old were deeply anesthetized with sodium pentobarbitol (200mg/kg; i.p.; VetOne) and transcardially perfused with 15mL ice-cold NMDG-artificial cerebro-spinal fluid (aCSF), continuously bubbled with carbogen (95% O_2_ + 5% CO_2_), and containing (in mM): 92 NMDG, 2.5 KCl, 1.25 NaH_2_PO_4_, 30 NaHCO_3_, 20 HEPES, 25 D-Glucose, 2 thiourea, 5 Na-ascorbate, 3 Na-pyruvate, 0.5 CaCl_2_, and 10 MgSO_4_. Brains were then extracted and 200 µm coronal slices were cut using a vibratome (Leica) containing ice-cold NMDG-aCSF. Slices were then transferred to a recovery chamber containing NMDG-aCSF at 31°C for 20-30 min. A 2M Na^+^ spike-in solution (116 mg/mL Na^+^ in NMDG-aCSF) was added to the recovery chamber in increasing volumes (from 250 µL to 1 mL) in 5 min increments for 25 min in order to achieve a controlled rate of reintroduction of Na^+^ into the chamber^3^. 5 min after the last Na^+^-spiking solution, slices were then transferred into room temperature HEPES-aCSF containing (in mM): 92 NaCl, 2.5 KCl, 1.25 NaH_2_PO_4_, 30 NaHCO_3_, 20 HEPES, 25 D-Glucose, 2 thiourea, 5 Na-ascorbate, 3 Na-pyruvate, 2 CaCl_2_, 2 MgSO_4_, and continuously bubbled in carbogen. After 30-45 min recovery, slices were transferred to a recording chamber continuously perfused with carbogenated aCSF (in mM: 124 NaCl, 2.5 KCl, 1.25 NaH_2_PO_4_, 24 NaHCO_3_, 5 HEPES, 12.5 D-Glucose, 2 CaCl_2_, 2 MgSO_4_) at a rate of 2-3 ml/min and maintained at 32°C by an in-line heater (Warner Instruments).

Patch pipettes (4.5-6.5 MΩ) were pulled from borosilicate glass (Kings Precision Glass) using a gravity puller (Narishige). For voltage-clamp experiments, pipettes were filled with a cesium-based internal solution containing (in mM): 130 D-Gluconic acid, 130 CsOH, 5 NaCl, 10 HEPES, 12 phosphocreatine, 3 Mg-ATP, 0.2 Na-GTP, 10 EGTA, at pH 7.25 and 285 mOsm^4^.

For current-clamp experiments, pipettes were filled with a K-methanesulfonate-based internal solution containing (in mM): 115 KMeSO_3_, 20 NaCl, 1.5 MgCl_2_, 0.1 EGTA, 5 HEPES, 2 Mg-ATP, 0.3 Na_2_-GTP, 10 Na_2_-Phosphocreatine, at pH 7.3 and 282 mOsm^5^. Epifluorescence was used to locate YFP-labeled VTA^GG^ terminals in LHb and DREADD-mCherry-labeled postsynaptic LHb neurons. Visually guided patch recordings were made using infrared differential interference contrast (IR_DIC) illumination (A1 Examiner, Zeiss). A light-emitting diode (UHP-LED460, Prizmatix) under computer control was used to flash blue light through the light path of the microscope to activate ChR2 terminals. Recordings were made using a Multiclamp 700B Amplifier (Axon Instruments), filtered at 2 kHz, and digitized at 10 kHz (Axon Digidata, Axon Instruments), and collected using pClamp v10 software (Molecular Devices). Capacitance and series resistance were electronically corrected before recordings, and series resistance monitored throughout recordings. Any cell in which series resistance changed >20% was discarded and excluded from analysis. Optogenetically-evoked excitatory-and inhibitory-postsynaptic currents (o-EPSC and o-IPSC) were voltage clamped at-60 mV and 0 mV, respectively, in whole cell configuration. A single 50 ms blue light pulse was applied every 45 s, and 10 light-evoked currents were averaged per neuron per condition. Picrotoxin (Sigma) was diluted in aCSF for 10 µM bath application. AMPA (alpha-amina-3-hydroxy-5-methyl-4-isoxazolepropionic acid) receptors were blocked using 6,7-dinitroquinoxaline-2,3-dione (DNQX, Sigma) dissolved in dimethyl sulfoxide (DMSO, Sigma) and diluted in aCSF for 10µM bath application. Optogenetically-evoked postsynaptic potentials were current clamped at I=0 in whole cell configuration. A single 50 ms blue light pulse was applied every 45 s, and 10 light-evoked potentials were averaged per neuron per condition. Resting membrane voltage was measured in current clamp at 0 pA during whole cell configuration and throughout current clamp recordings. For DREADD experiments, clozapine-N-oxide (NIDA Drug Supply Program 030534) was dissolved in DMSO and diluted in aCSF for 10µM bath application.

### 2 nose-poke optogenetic self-stimulation procedure

Mice were food restricted to 85-90% of baseline body weight 1-2 days prior to and during testing to increase baseline responding. During testing, mice were tethered to a 50 cm bilateral patch cord attached to an optical rotary joint (Doric), connected through an FC connector to laser (473 nm, Shanghai Laser & Optics Century). Mice were placed in operant chambers (Med Associates) controlled by MedPC IV software. Each session began with turning on of the house light, the LED cue lights above each of the 2 nose ports, and playing of a 0.5s tone (2 kHz). Each nose port contained photobeams and were baited with one 20 mg sucrose pellet (BioServ F0071) prior to each session. Beam breaks into each nose port (‘nose poke’) triggered a 0.5 s tone and turned off the LED lights for 1 s. Beam breaks into the active nose port also delivered laser stimulation (1 s, 10 mW, 40 Hz, 5 ms pulse width) through a TTL-generator controlled by an Arduino board. Nose pokes that occurred during the 1 s laser stimulation were recorded but had no consequence. Each session lasted 45 min. For DREADD experiments, mice received an injection of vehicle (0.5% DMSO in saline; i.p., 10 mL/kg volume) 15 min prior to sessions on days 1-3, and an injection of CNO (3 mg/kg, i.p., 10 mL/kg volume) 15 min prior to sessions on days 4-6.

#### Open Field

Mice were placed into an open field apparatus (45 cm W x 45 cm D x 40 cm H) for 20-min sessions. CNO (i.p., 3 mg/kg, 10 mL/kg volume) or vehicle (0.5% DMSO, i.p., 10 mL/kg volume) was injected 15 min prior to the sessions, with order of drug session counterbalanced between mice. CNO and vehicle sessions were separated by 48 hours. Distance travelled was collected using AnyMaze software attached to overhead cameras.

#### DREADD-mediated c-Fos induction

After recovery from DREADD or mCherry viral injection into LHb (5 weeks), mice were handled for 3 days and then habituated to a testing room ∼2 hr prior to injection. Mice were injected with CNO (3 mg/kg, i.p., 10 mL/kg volume) and transcardially perfused (as described above) 75 min later. Brains were extracted and processed as described above, and immunostained for c-Fos expression (as described above). Images of bilateral LHb were acquired at 10x magnification using a Zeiss Axio Observer. Boundaries of LHb were drawn and the total amount of Fos-expressing neurons were counted using ImageJ software.

#### Fiber photometry recordings

For all fiber photometry experiments, mice were placed in a plastic transfer cage (17 cm W x 30 cm D x 25.4 cm H) with an open top and tethered to a 100 cm-long patch cord (400 µm core, NA 0.48, Doric) attached to a pigtailed fiber optic rotary joint (Doric) connected through an FC connector to dichroic mirrors (minicube, Doric), and then to an LED driver (Doric) and red laser (635 nm, Shanghai Laser & Optics Century). An isosbestic channel (405 nm) was used to control for movement artifacts while 465 nm was used to stimulate GCaMP, and 635 nm laser was used to optogenetically excite VTA^GG^ terminals. Fluorescence was transmitted through a femtowatt (Newport, Doric), digitized at 1017 Hz, recorded by a real-time signal processor (RZ5P, TDT), and processed by Synapse software (TDT). Event timestamps were integrated by TTL inputs from AnyMaze. During each session, laser stimulation (635 nm) was delivered at a given frequency every 20 s, and in alternating durations such that each session delivered 10 trials of each duration (0.5 s, 1 s, 2 s, and 5s). Mice were tested at one frequency per session (5 Hz, 10 Hz, 20 Hz, and 40 Hz) (∼ 15 min per session), receiving 2 different frequency sessions per day such that photometry recordings lasted no more than ∼30-40 min a day. During fiber photometry sessions with DREADD activation, sessions began with 10 trials of laser stimulations separated by 20 s, alternating between frequencies and duration (10 Hz 1 s, 10 Hz 5s, 40 Hz 1 s, 40 Hz 5 s). After mice received 10 stimulations of each frequency/duration of laser, CNO (i.p., 3 mg/kg, 10 mL/kg volume) or vehicle (0.5% DMSO, i.p., 10 mL/kg volume) was injected while mice were tethered and fluorescence was being monitored. Laser stimulation sessions identical to the baseline session began 15 min later. Order of CNO and vehicle sessions were counterbalanced across mice.

Analysis of the recorded calcium signals was performed using custom-written MATLAB scripts. Fluorescent signal in response to the isosbestic channel (405 nm) was fitted to the signal at 465 nm, and then subtracted from the 465 nm signal to create a delta-F/F (dF/F). dF/F was monitored over a baseline, pre-stimulus window of 5 s, followed by 10 s post-stimulus, and subsequently normalized to a z-score based on the mean and standard deviation of the 5 s baseline (dF/F). Area under the curve during the pre-stimulus and stimulus windows (0.5 s – 5 s, depending on laser stimulation duration) was then extracted per animal and averaged per group.

### QUANTIFICATION AND STATISTICAL ANALYSIS

Data were analyzed using t-tests (one sample, unpaired, or paired), 2-way repeated-measure ANOVAs followed by Bonferroni or Holm-Sidak post-hoc multiple comparisons, one-way ANOVAs, and Pearson correlation (GraphPad Prism v7). Ordinal data or data that was not normally distributed was analyzed using Mann-Whitney tests or Kruskal-Wallis tests followed by Dunn’s post-hoc multiple comparisons tests. All data are represented as mean ± standard error of the mean (SEM) and/or as individual points. All statistical details of each experiment, including statistical tests used and sample sizes, can be found in figure legends.

## Acknowledgments

This work was supported by grants from NIH (R01DA036612, F32MH122192, P30DA048736, and K99/R00MH130688) and Veterans Affairs (I01BX005782). We thank Elizabeth Eckenweiler and William S. Conrad for assistance with mouse colony breeding. We are also grateful to Richard Palmiter and Sekun Park for providing a viral vector.

## Author Contributions

Conceptualization, S.M.W., V.Z., and T.S.H.; methodology, S.M.W., N.G.H., V.Z., and T.S.H.; investigation, S.M.W., D.S.D., N.G.H., and L.O.; formal analysis, S.M.W., L.F., and T.S.H.; writing – original draft, S.M.W. and T.S.H.; writing – review and editing, S.M.W., L.S.Z., and T.S.H.; resources, L.S.Z.; funding acquisition, S.M.W. and T.S.H.

## Competing Interests Declaration

The authors declare no competing interests.

## Resource Availability

Supplementary Information is available for this paper.

Correspondence and requests for materials should be addressed to Thomas Hnasko and Shelley Warlow.

## Notes

### Competing Interest Statement

The authors have declared no competing interest.

## Main References

1. Morales, M. & Margolis, E. B. Ventral tegmental area: cellular heterogeneity, connectivity and behaviour. Nat. Rev. Neurosci. 18, 73–85 (2017).

2. Wallace, M. L. & Sabatini, B. L. Synaptic and circuit functions of multitransmitter neurons in the mammalian brain. Neuron 111, 2969–2983 (2023).

3. Conrad, W. S., Oriol, L., Kollman, G. J., Faget, L. & Hnasko, T. S. Proportion and distribution of neurotransmitter-defined cell types in the ventral tegmental area and substantia nigra pars compacta. Addict. Neurosci. 13, (2024).

4. Root, D. H. et al. Single rodent mesohabenular axons release glutamate and GABA. Nat. Neurosci. 17, 1543–1551 (2014).

5. Root, D. H. et al. Selective Brain Distribution and Distinctive Synaptic Architecture of Dual Glutamatergic-GABAergic Neurons. Cell Rep. 23, 3465–3479 (2018).

6. Wang, H.-L., Qi, J., Zhang, S., Wang, H. & Morales, M. Rewarding effects of optical stimulation of ventral tegmental area glutamatergic neurons. J. Neurosci. 35, 15948–15954 (2015).

7. Warlow, S. M. et al. Mesoaccumbal glutamate neurons drive reward via glutamate release but aversion via dopamine co-release. Neuron 112, 488–499.e5 (2024).

8. Zell, V. et al. VTA Glutamate Neuron Activity Drives Positive Reinforcement Absent Dopamine Co-release. Neuron 107, 864–873.e4 (2020).

9. Yoo, J. H. et al. Ventral tegmental area glutamate neurons co-release GABA and promote positive reinforcement. Nat. Commun. 7, 13697 (2016).

10. Baker, P. M. et al. The Lateral Habenula Circuitry: Reward Processing and Cognitive Control. J. Neurosci. Off. J. Soc. Neurosci. 36, 11482–11488 (2016).

11. Matsumoto, M. & Hikosaka, O. Lateral habenula as a source of negative reward signals in dopamine neurons. Nature 447, 1111–1115 (2007).

12. Wang, D. et al. Learning shapes the aversion and reward responses of lateral habenula neurons. ELife 6, (2017).

13. Friedman, A. et al. Electrical stimulation of the lateral habenula produces enduring inhibitory effect on cocaine seeking behavior. Neuropharmacology 59, 452–459 (2010).

14. Stamatakis, A. M. et al. A unique population of ventral tegmental area neurons inhibits the lateral habenula to promote reward. Neuron 80, 1039–1053 (2013).

15. Brown, P. L. et al. Habenula-Induced Inhibition of Midbrain Dopamine Neurons Is Diminished by Lesions of the Rostromedial Tegmental Nucleus. J. Neurosci. 37, 217–225 (2017).

16. Lammel, S. et al. Input-specific control of reward and aversion in the ventral tegmental area. Nature 491, 212–217 (2012).

17. Browne, C. A., Hammack, R. & Lucki, I. Dysregulation of the lateral habenula in major depressive disorder. Front. Synaptic Neurosci. 10, 46 (2018).

18. Lecca, S., Meye, F. J. & Mameli, M. The lateral habenula in addiction and depression: an anatomical, synaptic and behavioral overview. Eur. J. Neurosci. 39, 1170–1178 (2014).

19. Loonen, A. J. M., Kupka, R. W. & Ivanova, S. A. Circuits regulating pleasure and happiness in bipolar disorder. Front. Neural Circuits 11, 35 (2017).

20. Kim, S., Wallace, M. L., El-Rifai, M., Knudsen, A. R. & Sabatini, B. L. Co-packaging of opposing neurotransmitters in individual synaptic vesicles in the central nervous system. Neuron 110, 1371–1384.e7 (2022).

21. Xu, J. et al. Intersectional mapping of multi-transmitter neurons and other cell types in the brain. Cell Rep. 40, 111036 (2022).

22. Seal, R. P. & Edwards, R. H. Functional implications of neurotransmitter co-release: glutamate and GABA share the load. Curr. Opin. Pharmacol. 6, 114–119 (2006).

23. Vaaga, C. E., Borisovska, M. & Westbrook, G. L. Dual-transmitter neurons: functional implications of co-release and co-transmission. Curr. Opin. Neurobiol. 29, 25–32 (2014).

24. Lammel, S. et al. Diversity of transgenic mouse models for selective targeting of midbrain dopamine neurons. Neuron 85, 429–438 (2015).

25. Brinschwitz, K. et al. Glutamatergic axons from the lateral habenula mainly terminate on GABAergic neurons of the ventral midbrain. Neuroscience 168, 463–476 (2010).

26. Stamatakis, A. M. et al. Lateral hypothalamic area glutamatergic neurons and their projections to the lateral habenula regulate feeding and reward. J. Neurosci. 36, 302–311 (2016).

27. Congiu, M. et al. Plasticity of neuronal dynamics in the lateral habenula for cue-punishment associative learning. Mol. Psychiatry 28, 5118–5127 (2023).

28. Lecca, S. et al. Aversive stimuli drive hypothalamus-to-habenula excitation to promote escape behavior. eLife 6, e30697 (2017).

29. Yang, Y. et al. Ketamine blocks bursting in the lateral habenula to rapidly relieve depression. Nature 554, 317–322 (2018).

30. Hnasko, T. S. & Edwards, R. H. Neurotransmitter corelease: mechanism and physiological role. Annu. Rev. Physiol. 74, 225–243 (2012).

31. Root, D. H., Mejias-Aponte, C. A., Qi, J. & Morales, M. Role of glutamatergic projections from ventral tegmental area to lateral habenula in aversive conditioning. J. Neurosci. 34, 13906–13910 (2014).

32. Cerniauskas, I. et al. Chronic Stress Induces Activity, Synaptic, and Transcriptional Remodeling of the Lateral Habenula Associated with Deficits in Motivated Behaviors. Neuron 104, 899–915.e8 (2019).

33. Gill, M. J., Ghee, S. M., Harper, S. M. & See, R. E. Inactivation of the lateral habenula reduces anxiogenic behavior and cocaine seeking under conditions of heightened stress. Pharmacol. Biochem. Behav. 111, 24–29 (2013).

34. Zhang, H. et al. Dorsal raphe projection inhibits the excitatory inputs on lateral habenula and alleviates depressive behaviors in rats. Brain Struct. Funct. 223, 2243–2258 (2018).

35. Lalive, A. L., Nuno-Perez, A., Tchenio, A. & Mameli, M. Mild stress accumulation limits GABAergic synaptic plasticity in the lateral habenula. Eur. J. Neurosci. (2021) doi:10.1111/ejn.15581.

36. Meye, F. J. et al. Shifted pallidal co-release of GABA and glutamate in habenula drives cocaine withdrawal and relapse. Nat. Neurosci. 19, 1019–1024 (2016).

37. Tchenio, A., Lecca, S., Valentinova, K. & Mameli, M. Limiting habenular hyperactivity ameliorates maternal separation-driven depressive-like symptoms. Nat. Commun. 8, 1135 (2017).

38. Shabel, S. J., Proulx, C. D., Piriz, J. & Malinow, R. Mood regulation. GABA/glutamate co-release controls habenula output and is modified by antidepressant treatment. Science 345, 1494–1498 (2014).

39. Destexhe, A., Rudolph, M. & Paré, D. The high-conductance state of neocortical neurons in vivo. Nat. Rev. Neurosci. 4, 739–751 (2003).

40. Gulledge, A. T., Kampa, B. M. & Stuart, G. J. Synaptic integration in dendritic trees. J. Neurobiol. 64, 75–90 (2005).

41. Poller, W. C. et al. Lateral habenular neurons projecting to reward-processing monoaminergic nuclei express hyperpolarization-activated cyclic nucleotid-gated cation channels. Neuroscience 193, 205–216 (2011).

42. Stuart, G. J. & Spruston, N. Dendritic integration: 60 years of progress. Nat. Neurosci. 18, 1713–1721 (2015).

43. Hirai, H. et al. Distinct release properties of glutamate/GABA co-transmission serve as a frequency-dependent filtering of supramammillary inputs. eLife 13, RP99711 (2024).

44. Li, S. et al. Synaptic sign switching mediates online dopamine updates. BioRxiv Prepr. Serv. Biol. 2025.07.23.666367 (2025) doi:10.1101/2025.07.23.666367.

## Methods References

1. Hunker, A. C. & others. Conditional single vector crispr/sacas9 viruses for efficient mutagenesis in the adult mouse nervous system. Cell Rep 30, 4303–4316.e6 (2020).

2. Grimm, D., Kern, A., Rittner, K. & Kleinschmidt, J. A. Novel tools for production and purification of recombinant adenoassociated virus vectors. Hum Gene Ther 9, 2745–2760 (1998).

3. Ting, J. T. & others. Preparation of Acute Brain Slices Using an Optimized N-Methyl-D-Glucamine Protective Recovery Method. (J. Vis. Exp, 2018). doi:10.3791/53825.

4. Franks, K. M. & Isaacson, J. S. Synapse-specific downregulation of NMDA receptors by early experience: a critical period for plasticity of sensory input to olfactory cortex. Neuron 47, 101–114 (2005).

5. Kramer, P. F. & Williams, J. T. Cocaine Decreases Metabotropic Glutamate Receptor mGluR1 Currents in Dopamine Neurons by Activating mGluR5. Neuropsychopharmacology 40, 2418–2424 (2015).

